# Holographic deep learning for rapid optical screening of anthrax spores

**DOI:** 10.1101/109108

**Authors:** YoungJu Jo, Sangjin Park, JaeHwang Jung, Jonghee Yoon, Hosung Joo, Min-hyeok Kim, Suk-Jo Kang, Myung Chul Choi, Sang Yup Lee, YongKeun Park

## Abstract

Establishing early warning systems for anthrax attacks is crucial in biodefense. Here we present an optical method for rapid screening of *Bacillus anthracis* spores through the synergistic application of holographic microscopy and deep learning. A deep convolutional neural network is designed to classify holographic images of unlabeled living cells. After training, the network outperforms previous techniques in all accuracy measures, achieving single-spore sensitivity and sub-genus specificity. The unique ‘representation learning’ capability of deep learning enables direct training from *raw images* instead of manually extracted features. The method automatically recognizes key biological traits encoded in the images and exploits them as fingerprints. This remarkable learning ability makes the proposed method readily applicable to classifying various single cells in addition to *B. anthracis*, as demonstrated for the diagnosis of *Listeria monocytogenes*, without any modification. We believe that our strategy will make holographic microscopy more accessible to medical doctors and biomedical scientists for easy, rapid, and accurate diagnosis of pathogens, and facilitate exciting new applications.

## Introduction

*Bacillus anthracis*, a gram-positive spore-forming bacterium causing the disease anthrax, is one of the most destructive biological weapons that is prone to be abused for bioterrorism (1). It is thus crucial to rapidly detect and identify anthrax spores for biodefense (2). Various biological, chemical, and optical fingerprinting methods have been studied to accelerate diagnosis of *B. anthracis* (3-5). Conventional culture-based methods take days and are often inaccurate. PCR-based methods provide species-level specificity but still take hours and require heavy instrumentation with skilled personnel to operate the system (4). Photoluminescence and surface-enhanced Raman scattering methods take only minutes, but require labeling with exogenous agents and cannot discriminate *B. anthracis* from other *Bacillus* species which are ubiquitous in nature (5). More importantly, most of these methods are limited by the detection sensitivity of at least thousands of bacterial cells; thus, their applications in practical settings such as aerosolized spores require sample amplification processes that significantly limit the detection speed.

Recent developments of optical methods based on holographic microscopy combined with machine learning, which enables rapid and label-free identification of *single* cells, could be an important step to address the anthrax issue (6-11). Holographic microscopy (12), or quantitative phase imaging (QPI) in a broader sense, measures optical field images (i.e., nanometer-scale distortions of wavefronts passing through a sample) using laser-based interferometry. In addition to the amplitude images available from conventional intensity-based microscopy techniques, holographic microscopy *quantitatively* measures the optical phase delay maps dictated by the refractive index (RI) distribution of a sample (12). Because the endogenous RI distribution in a cell is strongly related to the structural and biochemical characteristics (13) of the target classes (e.g., species or cell types), the measured field images of single cells and the corresponding class labels are passed to data-driven machine learning algorithms for systematic discovery of class-specific fingerprints encoded in the images. These approaches can be combined with flow cytometry and/or bioaerosol collection systems to achieve ultrafast identification of unlabeled cells and pathogens (9, 10). However, none of these methods achieved *sub-genus specificity* required for discriminating *B. anthracis* from other *Bacillus* species ubiquitous in nature.

Here we present a next-generation holographic screening method by adopting ‘deep learning’, a state-of-the-art machine learning technique based on deep multi-layered neural networks (14, 15), to holographic microscopy. We designed a deep convolutional neural network (CNN) HoloConvNet specialized in the classification of holographic images of living cells. After training with quantitative phase images of individual *Bacillus* spores, the network identified new anthrax spores with single-spore sensitivity and sub-genus specificity. Its remarkable learning ability enables direct training from *raw images* by automatically recognizing key biological traits encoded in the images, and presents outstanding accuracy that outperforms previous approaches in all accuracy measures. As demonstrated below, this method is readily applicable to classification of various single cells in addition to *B. anthracis* without any modification.

## Results

### The holographic deep learning framework

The overall framework of our method is shown in Fig. 1. We used quantitative phase imaging unit (QPIU), a cost-effective palm-sized module that converts a conventional microscope into a holographic microscope (16), for phase imaging of individual *Bacillus* spores in an isolated biosafety level 3 (BSL-3) laboratory at the Agency for Defense Development, Korea. It is attached to the output port of an existing bright-field microscope to form a common-path interferometry for optical field imaging (Fig. 1 *A-C*; see *Materials and Methods*). After imaging *B. anthracis* and four different *Bacillus* species with various levels of phylogenetic relatedness (see Supplementary Note 1), we trained our deep neural network named HoloConvNet as a species classifier using the phase images of individual spores and the corresponding species labels (training set). The learnable parameters of the deep neural network were iteratively adjusted by the error backpropagation algorithm(14, 15) (Fig. 1D). The performance of the trained HoloConvNet was tested by taking new images (test set), which were never seen before by the network, as the input to the network (Fig. 1E). The machine-predicted species labels were compared with the true classes to estimate identification accuracy.

**Fig. 1.**
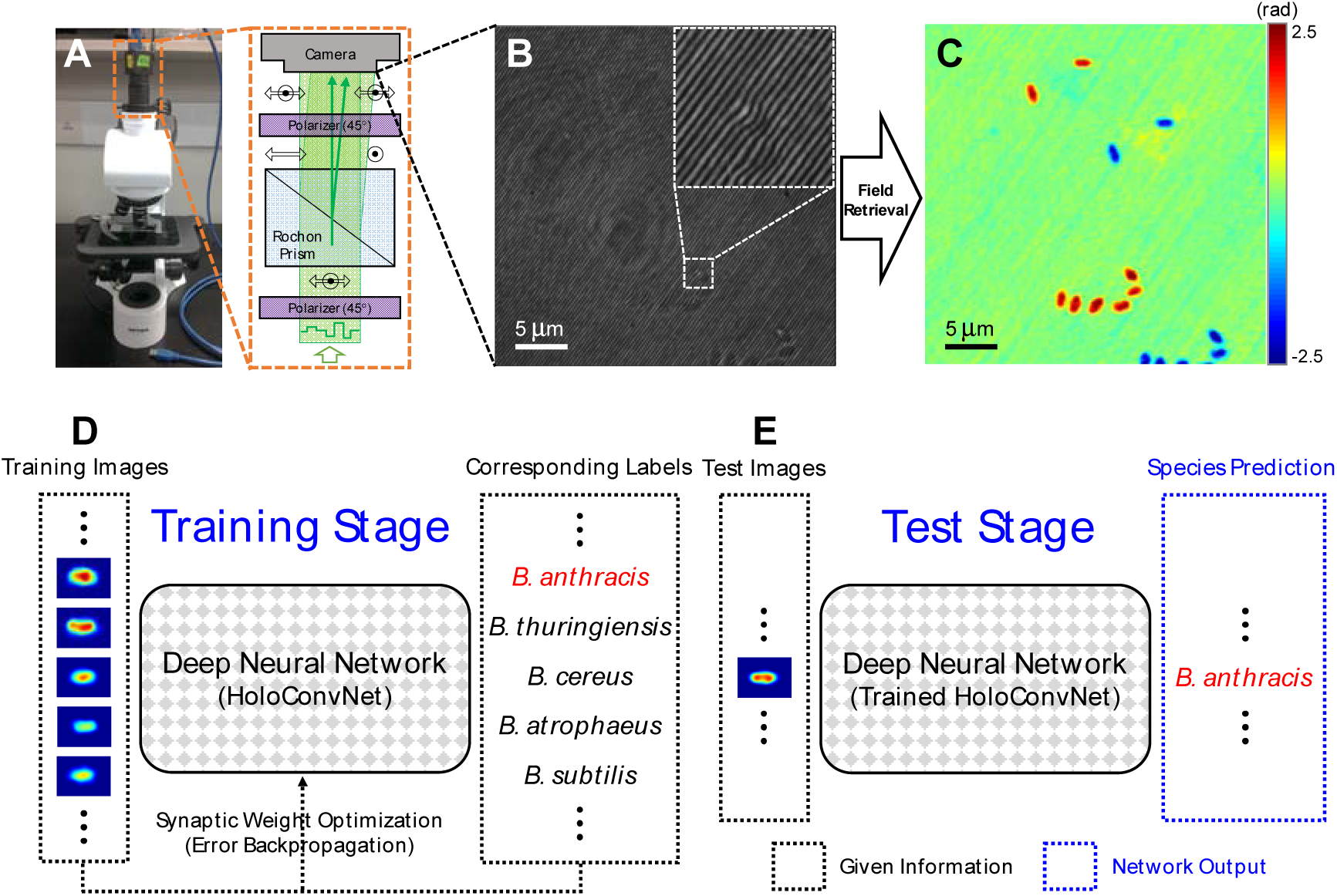
Holographic deep learning framework for screening of anthrax spores. (*A*) Schematic diagram of QPIUfor holographic imaging of individual *Bacillus* spores. (*B*) An interferogram formed by spatial modulation. It encodes quantitative phase images of individual spores as shown in (*C*). (*D*) The measured phase images from multiple *Bacillus* species are used to train a deep neural network using the error backpropagation algorithm. (*E*) The trained network accurately predicts the corresponding species when independently-measured phase images are shown.

The *quantitative* nature of holographic microscopy captures subcellular phase delay distribution which could be exploited by machine learning algorithms to extract fingerprint information (8, 13). On the other hand, conventional techniques (e.g., phase contrast microscopy) provide rough morphological information only (Fig. S1). Simple morphological parameters such as spore size (Fig. S2) are not enough for species discrimination due to high genetic similarities and large cell-to-cell variations (8, 16).

The endogenous RI distribution of *Bacillus* spores, which dictates the sample-induced phase delay imaged by QPIU, is strongly related to specific characteristics of each species (8, 13). However, because this relation is often *indirect*, it should be *approximated* using supervised learning. The precision of this function approximation obviously dominates the performance of the trained classifiers. Deep neural networks are *universal approximators* for virtually any arbitrarily nonlinear functions (15), while conventional machine learning techniques mostly rely on linear or only slightly nonlinear decision boundaries (8).

The network architecture of HoloConvNet is illustrated in Fig. 2 (and Table S1). A phase image of a single spore is processed by multiple layers of convolution, nonlinearity, and pooling operations, and then finally receives scored class labels through fully-connected layers. The network makes its prediction by selecting the final-layer neuron with the strongest activation. The key functional block of this process is a convolutional layer followed by nonlinearity:

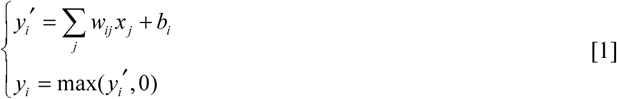

where *x* and *y* are input and output vectors, and *w* and *b* are synaptic weights and biases, respectively. Equation 1 emulates integration of synaptic inputs by a biological neuron (14, 15) that fires only when the net input exceeds a certain threshold (more precisely, a population of neurons with an output firing rate modeled by a rectified linear unit (ReLU)). Note that the entire processing by the network from images to class labels is a *nonlinear* mapping which corresponds to the approximating function explained above.

**Fig. 2.**
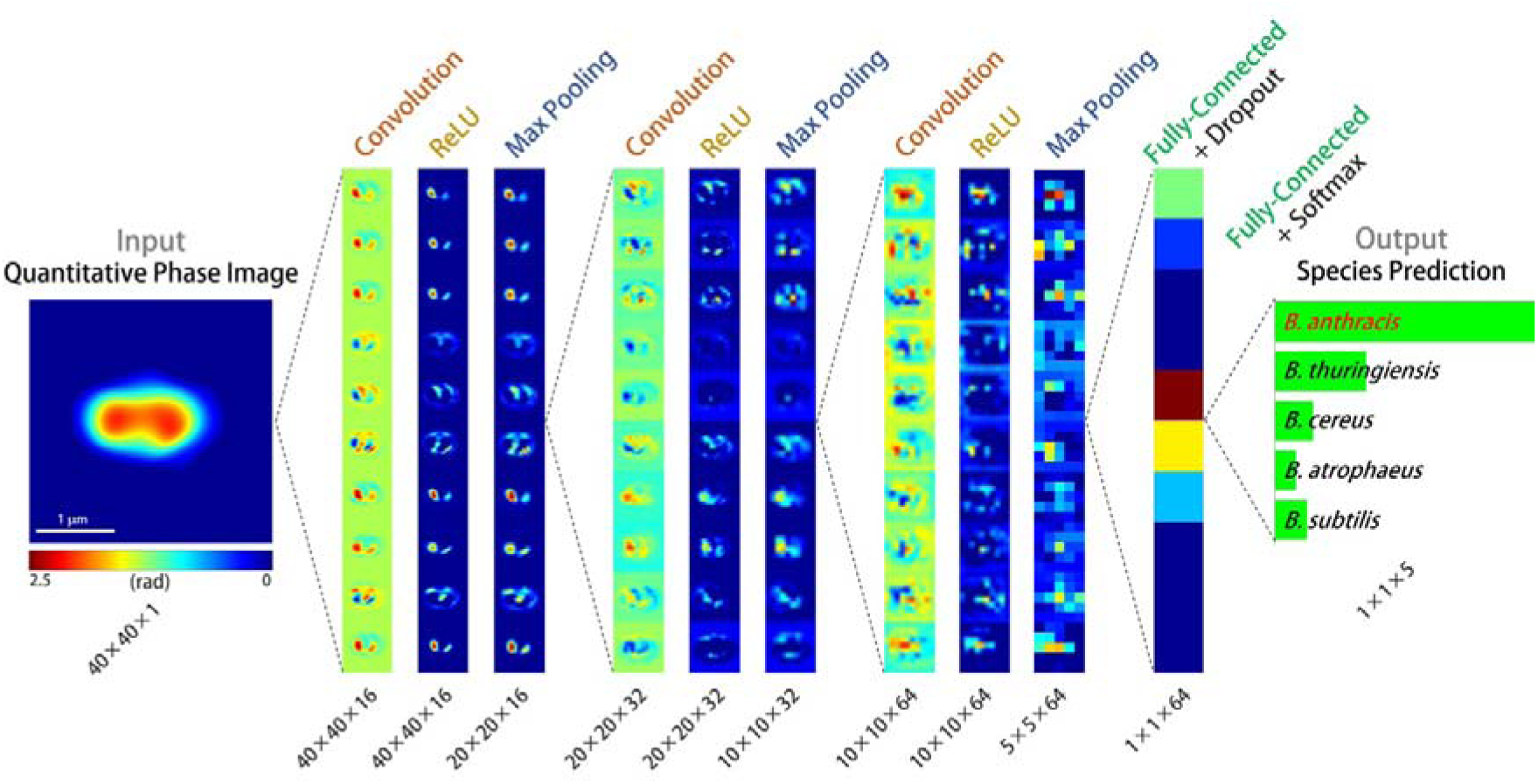
Architecture of HoloConvNet. When a phase image of an individual spore is taken as the input, thenetwork first processes the images through 3 rounds of convolution, ReLU nonlinearity, and max pooling layers. Then two fully-connected (and ReLU) layers follow; the first is the last hidden layer under dropout regularization, and the second is the output layer with the class scores. These scores are used to calculate the loss function and to make species predictions in the training and test stages, respectively. Only 10 two-dimensional activation maps per layer are presented with layer-wise scaling for visualization (see Table S1 for detailed architecture).

Training a deep neural network is essentially a large-scale nonlinear optimization of the synaptic weights (and biases) which govern the network behaviour. The large number of the learnable parameters makes training process extremely difficult. However, CNNs such as HoloConvNet have dramatically smaller number of parameters (14, 17) by employing localized and shared receptive field structures inspired by physiological visual processing (see Supplementary Note 2). Thus the network can be trained using the error back propagation algorithm that minimizes the mismatch between the machine-predicted and true labels (see *Materials and Methods*). HoloConvNet efficiently converges to a hierarchical *representation* of the images that gradually transforms the data space in which the classes are easily separable. This property is called the ‘representation learning’ capability of deep learning (14) and enables direct training from *raw images*.

### Performance

The performance of HoloConvNet is shown in Fig. 3. A well-trained neural network reflects the general relations between the input and output data, so that it accurately predicts the class labels of new images (*generalization* property). The multiclass identification performance of the network for the five *Bacillus* species, trained with 5 class labels representing individual species, is shown in Fig. 3A. HoloConvNet clearly identifies *B. anthracis* spores from the other four species with high sensitivity and specificity. Since diagnosing anthrax spores from other species is our prime objective, the network was next trained with binary class labels (anthrax versus non-anthrax). Using this method, the performance could be enhanced (Fig. 3B) by letting the optimization process focus on the characteristics distinguishing *B. anthracis* from others. When the problem was relaxed by excluding the two *Bacillus cereus* group species (note that these species are rare, while *Bacillus subtilis* is ubiquitous in nature; see Supplementary Note 1), HoloConvNet achieved a remarkable accuracy of 95.3% (Fig. 3C).

**Fig. 3.**
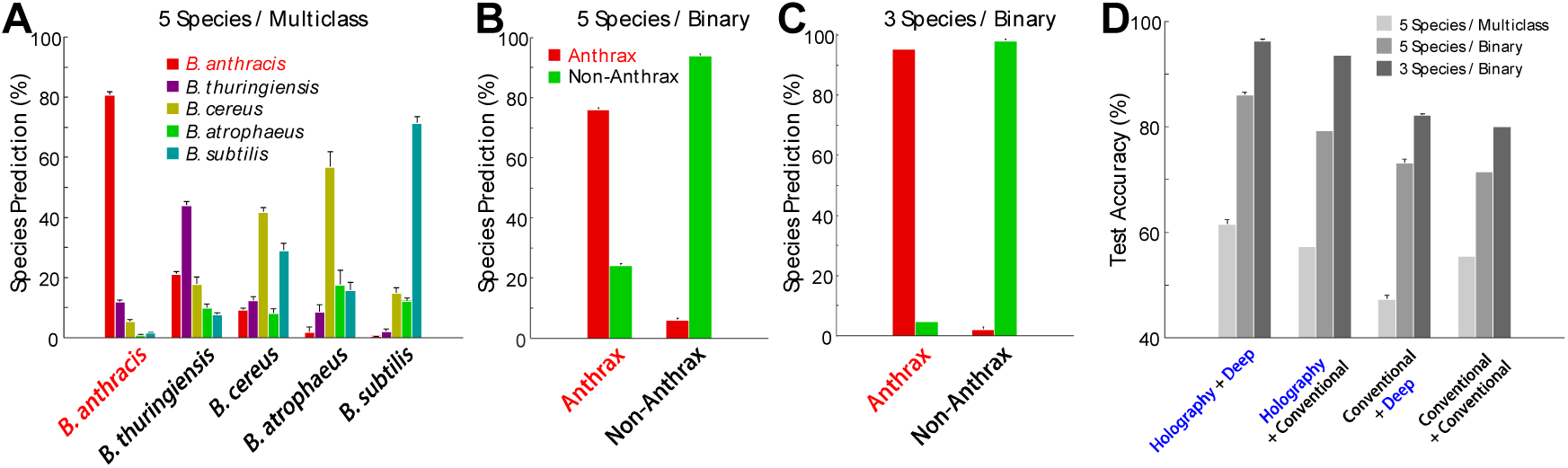
Performance of HoloConvNet. (*A*-*C*) The test images were used to measure the performance of (*A*)multiclass classification of 5 *Bacillus* species; (*B*) binary classification of *B. anthracis* and the other 4 species; (*C*) binary classification of *B. anthracis* and 2 non-member species of the *B. cereus* group. (*D*) The performance of the proposed method is compared to previous techniques (see the main text). Holographic microscopy and deep learning significantly improve the performance in all cases.

The performance of our method was compared with those of several previous techniques (Fig. 3D): holographic microscopy with conventional machine learning (8), conventional microscopy with deep learning (training HoloConvNet with binary morphology images; see *Materials and Methods*), and conventional microscopy with conventional machine learning (linear discriminant analysis with the morphological parameters in Fig. S2). HoloConvNet outperformed the previous methods in all accuracy measures, which clearly demonstrates the great potential of deep-learning-based holographic screening of anthrax spores in realistic settings.

### Representation learning

Representation learning by HoloConvNet, the fundamental improvement developed in this study, was further examined (Fig. 4). The network transforms the images into a representation in which the data points are linearly separable because a single layer of neurons is a linear classifier (15). We applied t-distributed stochastic neighbor embedding (t-SNE), a high-dimensional data visualization technique (18), to the activation of individual neurons in the last hidden layer (Fig. 4 *A-C*; see *Materials and Methods*). The nice separation observed indicates the great ability of HoloConvNet to learn the optimal representation of phase images without any pre-designed features required by conventional machine learning techniques. The different degrees of separation in the three cases explain the different identification performance. Additionally, the relative distances between the species clusters shown in Fig. 4A are consistent with the phylogenetic relationship (see Supplementary Note 1); here it should be noted that the relationship was independently discovered by HoloConvNet through training.

**Fig. 4.**
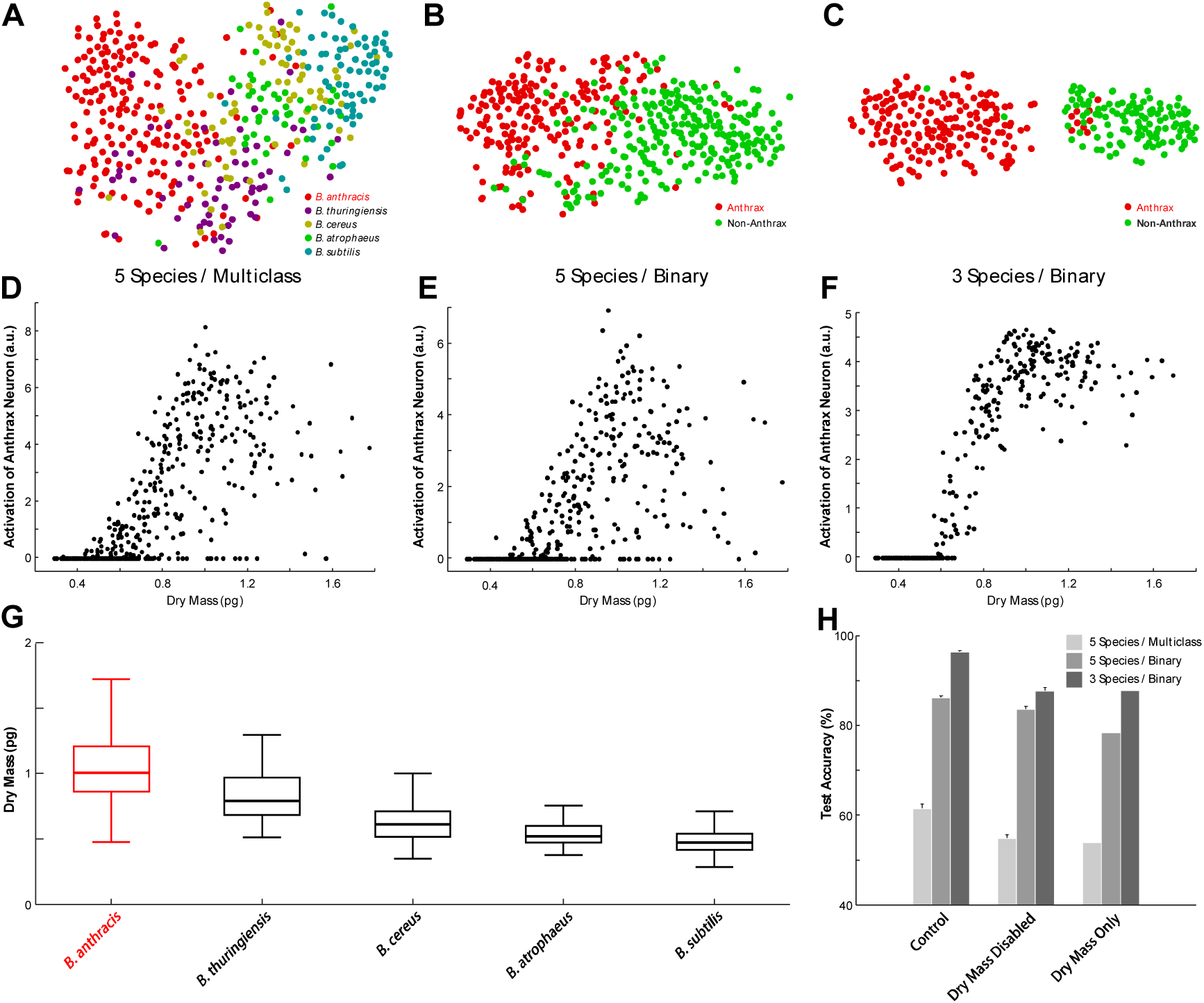
Representation learning by HoloConvNet. The inter-species difference in cellular dry mass is automatically recognized and used for screening of anthrax spores. (*A-C*) t-SNE visualization of the CNN codes at the last hidden layer, which shows the representation learning capability of HoloConvNet (see the main text). (*D*-*F*) The activation of the ‘anthrax neuron’ at the output layer shows a strong correlation with dry mass. (*G*), Dry mass of individual *Bacillus* spores calculated from the quantitative phase images. (*H*) Computationally disabling the dry mass information significantly impairs the performance of HoloConvNet. Dry mass alone is not enough for full performance as well.

The outstanding performance of the proposed method raises a question: what are the *key biological traits* that are measured and exploited for the identification of anthrax spores? We speculated that *cellular dry mass* (19), the mass of non-aqueous cellular components, is one of the most important traits. This hypothesis is based on the domain knowledge that there exists an additional outermost structure, called the exosporium, in the *B. cereus* group spores but not in the remaining two species (20). It was reasoned that structural distinction might result in an inter-species difference of dry mass which is inherently measured by holographic imaging with femtogram-level sensitivity (see *Materials and Methods*). Indeed, a strong positive correlation was found between dry mass and activation of the ‘anthrax neuron’ at the output layer (Fig. 4 *D-F*). This observation makes sense if the mean dry mass of *B. anthracis* is the heaviest among the five species, which turns out to be true (Fig. 4G). As expected, *B. anthracis* is slightly heavier than the other two *B. cereus* group species, and the remaining two species lacking exosporium have considerably lighter dry mass. The subtle difference within the *B. cereus* group might be due to species-dependent compositions and nanostructures of exosporium (21), while their contribution to dry mass should be confirmed by additional investigations. It was noted that the same order relation of dry mass was observed in all independent measurements (Fig. S3), and the overall range of measured dry mass is consistent with previous studies (16, 22).

To confirm the causality between dry mass and species prediction by HoloConvNet, a computational disabling strategy was employed. Detrimental effects on the performance were observed by computationally normalizing the phase images to remove the dry mass information. As shown in Fig. 4H, the network trained and tested with normalized phase images shows a significantly impaired performance, supporting the key role of dry mass. However, it does not mean that it is the sole information extracted by the network; the performance of a single-feature linear discriminant classifier based solely on dry mass was also significantly worse. This suggests that other traits such as spatial distribution of subcellular components in the spores play roles in screening. From these observations, it can be concluded that the inter-species difference of dry mass is recognized and exploited through representation learning by HoloConvNet. Here, it should be emphasized that we never taught the network on how to calculate dry mass from phase images. On the other hand, a conventional machine learning algorithm cannot make use of dry mass unless it is manually selected by a researcher.

Finally, the *generality* of our method expected from the outstanding learning abilities was investigated. As a proof-of-concept example, HoloConvNet was trained for diagnosing the pathogen *L. monocytogenes*, the causative agent of listeriosis which is often fatal to neonates and the elderly (23), from five different *Listeria* species. The diagnostic accuracy was surprisingly high, showing higher than 85%. The architecture and learning rules were identical to those used for the diagnosis of *Bacillus* species. It is also noted that *L. monocytogenes* is not the species with the heaviest dry mass in this case (Fig. S4). These results suggest that the holographic deep learning framework reported here has immediate and wide applicability in contrast to problem-specific conventional machine learning approaches.

## Discussion

We proposed and experimentally demonstrated a novel method for screening of anthrax spores by combining holographic microscopy and deep learning for the first time. The new strategy enables rapid label-free identification of individual anthrax spores with *sub-genus* specificity extending our previous *inter-genus* bacterial fingerprinting method based on conventional machine learning (8). In addition to the superior performance due to the extreme flexibility of deep neural networks, the transition from classical machine learning to deep learning fundamentally transforms holographic single-cell identification techniques by acquiring the *representation learning* capability. HoloConvNet automatically recognizes and then uses key biological characteristics which are species-dependent (e.g., dry mass in the anthrax problem) simply from *raw* images. Additionally, the present method can be readily extended to other single-cell classification problems, such as the diagnosis of *L. monocytogenes* demonstrated in this study, without any modification. Thus, our method eliminates the need to manually design and optimize features based on trial-and-error for individual problems.

The next steps beyond this proof-of-concept study to achieve practical ultrafast screening of anthrax spores are straightforward. Above all, the proposed method should be combined with flow cytometry (9, 10) and bioaerosol collection(24) systems to fully exploit the single-spore and label-free nature of the method. Then, a large amount of holographic imaging data from the resultant high-throughput device would be used to train HoloConvNet for more species and strains under various environmental conditions to assure stable field performance. The performance could be further improved by adopting multimodal QPI (e.g., spectral (25), polarimetric (26), or tomographic (27) images as the stacked input to the network) to increase the amount of raw information investigated by the network.

Despite the fast and label-free nature of holographic microscopy, the limited chemical specificity has left this tool overshadowed by fluorescence microscopy. Specific domain knowledge (e.g., homogeneity of hemoglobin concentration in red blood cells, high RI of lipid droplets in eukaryotic cells, etc.) has been required for effective use of the technique. The method proposed in this paper solves this difficulty by using the powerful learning abilities of deep neural networks. As we demonstrated in this study, now *intelligent* holographic microscopy can actively recognize and exploit the class-specific fingerprints, encoded in the raw images of various biological samples, without any prior knowledge. We believe that our strategy will make holographic microscopy more accessible to medical doctors and biomedical scientists for easy, rapid, and accurate diagnosis of pathogens, and facilitate exciting new applications.

## Materials and Methods

### Preparation of Bacillus spores

*Bacillus anthracis* Sterne (pXO1+ and pXO2-) was obtained from the Centers for Disease Control and Prevention, Korea (KCDC). *Bacillus thuringiensis* BGSC 4AJ1 was obtained from the Bacillus Genetic Stock Center (BGSC). *Bacillus cereus* ATCC 4342 was obtained from the American Type Culture Collection (ATCC). *Bacillus atrophaeus* KCCM 11314 was obtained from the Korean Culture Center for Microorganisms (KCCM). *Bacillus subtilis* 168 was obtained from the Korean Collection for Type Cultures (KCTC).

All experiments involving *B. anthracis* were conducted in a BSL-3 laboratory following the regulations in the Republic of Korea. Bacterial cells from frozen glycerol stocks were streaked onto Luria-Bertani (LB) agar plates and incubated at 30°C overnight. Next day, a single colony was inoculated into 5 mL of LB broth in a 50 mL CELLSTAR CELLreactor tube (Greiner Bio-One, Austria) and incubated at 30°C with shaking (200 rpm) for 8 hours. Then, 250 μL of the culture broth were transferred to 25 mL of GYS sporulation medium (28) in a 125 mL polycarbonate Erlenmeyer flask with a vent cap (Corning, NY) and incubated at 30°C with shaking (200 rpm) for 48 hours. After sporulation was completed, spores were harvested by centrifugation (5,420×g, 4°C) and washed four times with phosphate-buffered saline (PBS; Life Technologies, CA). Finally, the spores were suspended in 5 mL of PBS and stored at 4°C until use. Note that we prepared all the species with the same procedure.

A small volume (approximately 10 μL) of the bacterial solution was placed in an imaging chamber comprised of standard cover glasses (C024501, Matsunami Glass, Japan) and (optional) spacers with a thickness of 20-30 μm. Imaging was performed at room temperature after the spores settled down to the bottom and spread into a single layer. All *Bacillus* experiments were independently repeated three times.

### Holographic imaging

Because all anthrax experiments had to be conducted in a separate BSL-3 facility at the Agency for Defense Development, we used a compact and portable QPIU recently developed in our group(16), as the holographic imaging modality. It consists of two polarizers (LPVISE100-A, Thorlabs Inc., NJ) and a Rochon prism (#68-824, Edmund Optics Inc., NJ) inside an aluminum tube mounted in front of a CCD camera (FL3-U3-88S2C-C, PointGrey, Canada). Inserting the unit into the output port of a conventional bright-field microscope (B-382PLi-ALC, Optika, Italy) converts it into a holographic microscope. The light source for illumination was a diode laser (CPS532, λ = 532 nm, 4.5 mW, Thorlabs Inc., NJ), and the total magnification was ×100 determined by an objective lens (M-148, NA 1.25, oil-immersion, Optika, Italy). Acquisition time per interferogram was less than 20 ms, which could be reduced by many orders with high-intensity light sources and more sensitive cameras.

QPIU, shown in Fig. 1A, is a spatially-modulated self-reference interferometry. When the light passing through the sample encounters the unit, it becomes linearly polarized by the front polarizer. Then, the following Rochon polarizing prism divides the beam into two duplicated beams with slightly different propagation directions. Finally, the orthogonal polarization states of the divided beams become parallel by the rear polarizer. Thus, the two beams of parallel polarization generate an interference pattern at the overlapped region on the CCD plane. The linear polarizers before and after the prism are adjusted so that the interferogram has a high visibility (Fig. 1B). The quantitative phase information is retrieved (Fig. 1C) from the measured interferogram using a standard field-retrieval algorithm (29). The details on the principle of QPIU can be found elsewhere(16).

### Image analysis

All image analysis procedures were done with MATLAB (R2014b; MathWorks Inc., MA). The reconstructed phase images containing multiple spores were segmented by phase thresholding to be separated into images of single spores. The isolated spores were computationally aligned at the centres of square backgrounds for further analysis. The segmented regions were considered as the morphologies of individual spores that could be measured with conventional microscopy techniques such as phase contrast microscopy. The representative morphological parameters plotted in Fig. S2 were quantified with the *regionprops* function of MATLAB.

Calculation of the single-spore dry mass from phase images exploited the well-known proportionality between the optical phase delay and cellular dry mass (30). The total dry mass (*m*) can be calculated from the phase delay map 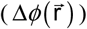 as follows:

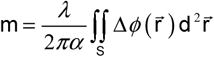

where *λ* is the illumination wavelength, *S* is the projection area of the cell surface, and *α* is the RI increment for non-aqueous molecules. Because the RI increment is known to be 0.18–0.21 mL/g for typical biological cells (30), we used *α* = 0.2 mL/g, and the results were consistent with those measured by other techniques (16, 22, 31). Note that we never explicitly taught the network about this relation.

### Deep learning

HoloConvNet is a CNN designed for the classification of holographic images of single cells. We implemented HoloConvNet using MatConvNet (32) framework (version 1.0 - beta 20) due to its simplicity and compatibility with our experimental data primarily processed with MATLAB. The final network architecture shown in Fig. 2 and Table S1 was carefully chosen after comparing several variations. Motivated by the recent trend for ‘small receptive fields and deep layers,’ the sizes of the receptive fields of the convolutional layers were chosen to be small (3-by-3), and thus, the total number of learnable parameters was relatively manageable (approximately 0.1 million). Because the architecture is substantially deep, we used the ReLU nonlinearity as the neuron model to avoid the vanishing gradient problem (33). Note that we used only phase images and not noisy amplitude images (due to the transparency and small sizes of the single spores) as the inputs to the network.

In addition to the traditional ‘weight decay’ regularization (15), several recent techniques were used to reduce overfitting. We used the ‘dropout’ technique, a regularization method based on efficient ensemble learning (34), for the last hidden layer with a dropout rate of 0.5. For further regularization and accelerated training speed, ‘batch normalization’ was done at every interface between a convolutional layer and the following ReLU layer (35). Due to the large number of learnable parameters, we used ‘data augmentation’ that enlarged the training dataset (17) by a factor of 128. This is simply done by generating the new labeled images by rotating the original images by random angles sampled from a zero-mean Gaussian distribution with a standard deviation of 10 degrees and by flipping the images with a probability of 0.5.

During the training stage, the learnable parameters were updated toward the direction minimizing the loss function using the concept of error backpropagation (14, 15). We used cross-entropy loss based on softmax function, which quantifies the mismatch between the machine-predicted and true labels, as the objective function to be minimized. By calculating the partial derivatives of the loss function with respect to the elements of the synaptic weight tensors using the chain rule, we could update the parameters in a stochastic gradient descent (SGD) scheme (14, 15). The learning rule was conventional SGD assisted with a momentum of 0.5 (note that using recent learning rules instead could further improve the performance), and the training batch size was 1024. The weights were initialized from a zero-mean Gaussian distribution with layer-wise scaling based on the input sizes (36). The biases were initialized with the constant 0. We used an equal learning rate for all layers, which was attenuated by a factor of 5 per five epochs (15). The hyperparameters were selected by cross-validation; the grid searching process with the initial learning rate and weight decay regularization strength resulted in values of 0.05 and 0.0005, respectively. We used one GPU (GeForce GTX 680, NVIDIA, CA) and CUDA Toolkit 7.5 (NVIDIA, CA), which increased the training speed typically by 5-10 fold. We note that it is possible with more computing resources to train multiple network models with different random initializations to compose a committee machine to further enhance the performance (17). Finally, the identification performance was estimated using separate test images which were never shown during the training stage. The error bars in Figs. 3 and 4 represent standard deviation calculated from 10 classification models with different random initializations.

The visualization of HoloConvNet codes was performed by the unsupervised dimensionality reduction technique t-SNE, which embeds high-dimensional data in a low-dimensional space while preserving the pairwise distances of the data points, implemented in MATLAB (18). The activation strengths of individual neurons at the last hidden layer by the test images were used as the raw variables. The parameters for the stochastic optimization for t-SNE were as follows: the perplexity was 30, and the dimension for initial principal component analysis was 30.

### The *Listeria* experiments

The six major bacterial species of the genus *Listeria*, *Listeria monocytogenes* (10403S), *Listeria grayi* (ATCC 19120), *Listeria innocua* (ATCC 33090), *Listeria ivanovii* (ATCC 19119), *Listeria seeligeri* (ATCC 35967), and *Listeria welshimeri* (ATCC 35897), were cultured in Brain-Heart Infusion media without antibiotics. After culturing overnight in a 37°C shaking incubator, the vegetative bacterial cells were washed and diluted with PBS based on the cultured concentration estimated by optical density measurements at 600 nm. The bacterial solution was placed and imaged in imaging chambers described above for the *Bacillus* experiments.

The holographic imaging of prepared samples was done with a Mach-Zehnder interferometry (12) with varying illumination angles to exploit the high-resolution synthetic aperture imaging technique (37). Optical field reconstruction and image processing protocols were identical to those of the *Bacillus* experiments.

The same network architecture and learning rule for training the original HoloConvNet (for *Bacillus* spores) were used to train the network for *Listeria*. The only preprocessing was to adjust the size of the input images to match the input dimension of HoloConvNet.

## Acknowledgements

We thank Minho Choi (KAIST) and SangHyun Kim (Samsung Electronics) for assisting the *Listeria* experiments; Kyeong Rok Choi (KAIST) and Seoyun Yum (UTSW) for biological insights; KyeoReh Lee, Kyoohyun Kim, and SangYun Lee (KAIST) for discussions; and Prof. Soo-Young Lee (KAIST) for inspiring lectures on neural networks. This work was supported by KAIST, ADD (ADD-14-70-06-10), National Research Foundation of Korea (2015R1A3A2066550, 2014K1A3A1A09063027, 2014M3C1A3052567), and Tomocube Inc. Y.-J.J. acknowledges support from KAIST Presidential Fellowship.

## Author Contributions

Y.-J.J. and Y.-K.P. conceived of the original idea for holographic deep learning. S.Y.L. and Y.-K.P. initiated and coordinated the project. Y.-J.J. and H.J. developed and analyzed HoloConvNet framework. J.-H.J. and J.Y. implemented QPIU and analyzed the holographic data. S.P. under the supervision of M.C.C. and S.Y.L. conducted the *Bacillus* experiments. Y.-J.J. and M.-H.K. under the supervision of S.-J.K. and Y.-K.P. performed the *Listeria* experiments. Y.-J.J., S.P., J.-H.J., J.Y., and Y.-K.P. wrote the initial version of the manuscript. All authors revised the manuscript.

## Competing Financial Interests

Y.-K.P. has financial interests in Tomocube Inc., a company that commercializes optical diffraction tomography and phase imaging instruments, and is one of the sponsors of this work.

## References

1. Brachman PS (2002) Bioterrorism: an update with a focus on anthrax. American Journal of Epidemiology 155(11):981–987.

2. WaltDR & FranzDR (2000) Peer Reviewed: Biological Warfare Detection. Analytical Chemistry 72(23):738A–746A.

3. KingD, LunaV, CannonsA, CattaniJ, & AmusoP (2003) Performance assessment of three commercial assays for direct detection of Bacillus anthracis spores. Journal of Clinical Microbiology 41(7):3454–3455.

4. HurtleW, et al. (2004) Detection of the Bacillus anthracis gyrA gene by using a minor groove binder probe. Journal of Clinical Microbiology 42(1):179–185.

5. ZhangX, YoungMA, LyandresO, & Van DuyneRP (2005) Rapid detection of an anthrax biomarker by surface-enhanced Raman spectroscopy. Journal of the American Chemical Society 127(12):4484–4489.

6. JavidiB, DaneshpanahM, & MoonI (2010) Three-Dimensional Holographic Imaging for Identification of Biological Micro/Nanoorganisms. IEEE Photonics Journal 2(2):256–259.

7. MoonI, DaneshpanahM, AnandA, & JavidiB (2011) Cell identification computational 3-D holographic microscopy. Optics and Photonics News 22(6):18–23.

8. JoY, et al. (2015) Label-free identification of individual bacteria using Fourier transform light scattering. Optics Express 23(12):15792–15805.

9. VercruysseD, et al. (2015) Three-part differential of unlabeled leukocytes with a compact lens-free imaging flow cytometer. Lab on a Chip 15(4):1123–1132.

10. Chen CL, et al. (2016) Deep Learning in Label-free Cell Classification. Scientific Reports 6:21471.

11. ParkHS, RinehartMT, WalzerKA, ChiJ-TA, & WaxA (2016) Automated Detection of P. falciparum Using Machine Learning Algorithms with Quantitative Phase Images of Unstained Cells. PLoS One 11(9):e0163045.

12. LeeK, et al. (2013) Quantitative phase imaging techniques for the study of cell pathophysiology: from principles to applications. Sensors 13(4):4170–4191.

13. LiuP, et al. (2016) Cell refractive index for cell biology and disease diagnosis: past, present and future. Lab on a Chip 16(4):634–644.

14. LeCunY, BengioY, & HintonG (2015) Deep learning. Nature 521(7553):436–444.

15. HaykinSS, HaykinSS, HaykinSS, & HaykinSS (2009) Neural networks and learning machines (Pearson).

16. JoY, et al. (2014) Angle-resolved light scattering of individual rod-shaped bacteria based on Fourier transform light scattering. Scientific reports 4:5090.

17. KrizhevskyA, SutskeverI, & HintonGE (2012) Imagenet classification with deep convolutional neural networks. Advances in Neural Information Processing Systems, pp 1097–1105.

18. Maaten Lvd & HintonG (2008) Visualizing data using t-SNE. Journal of Machine Learning Research 9(Nov):2579–2605.

19. PopescuG, et al. (2008) Optical imaging of cell mass and growth dynamics. American journal of physiology. Cell physiology 295(2):C538–544.

20. HenriquesAO & MoranJ, CharlesP (2007) Structure, assembly, and function of the spore surface layers. Annual Review of Microbiology 61:555–588.

21. Ball DA, et al. (2008) Structure of the exosporium and sublayers of spores of the Bacillus cereus family revealed by electron crystallography. Molecular Microbiology 68(4):947–958.

22. CarreraM, ZandomeniRO, & SagripantiJL (2008) Wet and dry density of Bacillus anthracis and other Bacillus species. Journal of Applied Microbiology 105(1):68–77.

23. LowJ & DonachieW (1997) A review of Listeria monocytogenes and listeriosis. The Veterinary Journal 153(1):9–29.

24. Després VR, et al. (2012) Primary biological aerosol particles in the atmosphere: a review. Tellus B 64.

25. JungJ-H, JangJ, & ParkY (2013) Spectro-refractometry of Individual Microscopic Objects Using Swept-Source Quantitative Phase Imaging. Analytical Chemistry 85(21):10519–10525.

26. KimY, JeongJ, JangJ, KimMW, & ParkY (2012) Polarization holographic microscopy for extracting spatio-temporally resolved Jones matrix. Optics Express 20(9):9948–9955.

27. KimK, et al. (2013) High-resolution three-dimensional imaging of red blood cells parasitized by Plasmodium falciparum and in situ hemozoin crystals using optical diffraction tomography. Journal of Biomedical Optics 19(1):011005–011005.

28. YoustenA & RogoffM (1969) Metabolism of Bacillus thuringiensis in relation to spore and crystal formation. Journal of Bacteriology 100(3):1229–1236.

29. DebnathSK & ParkY (2011) Real-time quantitative phase imaging with a spatial phase-shifting algorithm. Optics Letters 36(23):4677–4679.

30. BarerR (1952) Interference microscopy and mass determination. Nature 169(4296):366–367.

31. GodinM, et al. (2010) Using buoyant mass to measure the growth of single cells. Nature Methods 7(5):387–390.

32. VedaldiA & LencK (2015) Matconvnet: Convolutional neural networks for matlab. Proceedings of the 23rd ACM International Conference on Multimedia, (ACM), pp 689–692.

33. NairV & HintonGE (2010) Rectified linear units improve restricted boltzmann machines. Proceedings of the 27th International Conference on Machine Learning, pp 807–814.

34. SrivastavaN, HintonG, KrizhevskyA, SutskeverI, & SalakhutdinovR (2014) Dropout: A simple way to prevent neural networks from overfitting. Journal of Machine Learning Research 15(1):1929–1958.

35. IoffeS & SzegedyC (2015) Batch Normalization: Accelerating Deep Network Training by Reducing Internal Covariate Shift. Proceedings of The 32nd International Conference on Machine Learning, pp 448–456.

36. HeK, ZhangX, RenS, & SunJ (2015) Delving deep into rectifiers: Surpassing human-level performance on imagenet classification. Proceedings of the IEEE International Conference on Computer Vision, pp 1026–1034.

37. LeeK, et al. (2013) Synthetic Fourier transform light scattering. Optics Express 21(19):22453–22463.

